# The unapparent effect of climate forcing on urban amoebiasis in Mexico City

**DOI:** 10.1101/2021.01.17.427028

**Authors:** Andres Baeza, Mauricio Santos-Vega, Ana E. Escalante, Hallie Eakin

## Abstract

Despite advances in the provision of sanitation infrastructure in cities, diarrheal-related illnesses continue to be a global burden. In cities of the developing world, explanations for the persistence of diarrheal-related diseases revolve around the role of sanitation, poverty, and individual behavior. Less is known about the role of climate in forcing the inter-annual variability of transmission. We quantified the contribution of rainfall to the transmission of a diarrhoeal disease caused by the protozoan Entamoeba histolytica. We formulated a process-based model of the population dynamics of Entamoeba histolytica in 6 municipalities of Mexico City, and we reconstructed the interannual variability of the observed cases of amebiasis between 2005 and 2012. Using inference methods for non-linear systems, we estimated that 10% of the susceptible individuals were infected in early 2005 and 90% of those infections were causally related to the exposure to the environmental stage of the pathogen after rainfall events. The magnitude of these results underscores the importance of considering the interaction between climate forcing and water management as ecological determinants of health in megacities.

## 1 Introduction

Amoebiasis, a disease caused by the unicellular enteric protozoan *Entamoeba histolytica* (hereafter referred to as *Eh*), ranks third in protozoa-related mortalities (Sepulveda 1980; Esrey et al. 1985; Petri Jr et al. 2000; Haque et al. 2003; Chalmers 2014; Ortega and Verastegui 2018), causing 490,000 annual deaths, mostly children. Amoebiasis is also a leading factor in lost work and education opportunities in cities, exacerbating poverty, and inequality.

Cities of the developing world in which *Eh* is endemic are also affected by periodic environmental extremes such as flooding and drought. Extreme rainfall can overwhelm drainage infrastructure, exposing populations to contaminated water in which *Eh* thrives. Although there is an increasing concern over the risk of urban flooding under future climate change scenarios, the contribution of seasonal rainfall events to the magnitude of amoebiasis transmission in cities remains unclear (Singh et al. 2001; Drayna et al. 2010; Chou et al. 2010; Fletcher et al. 2012; Santos-Vega et al. 2016b).

Hypotheses on the important routes of transmission of *Eh* in cities revolve around the role of poverty, inadequate sanitation, and insufficient hygiene. Thus, without a vaccine, interventions to reduce its prevalence have centered on promoting individual behavioral changes through education (Sheth and Obrah 2004), rather than on reducing environmental risk through investments in infrastructure (Sanchez-Vega et al. 2006). Interventions to reduce environmental risk are often reactive and only triggered after episodic climate events, such as intense floods (Gibson et al. 1999).

Entamoeba histolytica relies on humans as primary (if not single) host(Othman et al. 2020). Exposure to *Eh* is initiated by the ingestion of water and food contaminated with environmentally resistant cysts. After an incubation period of 7 to 15 days, about 10-20% of exposed hosts present symptoms(Moonah et al. 2013). The symptoms of *Eh* are caused by the presence of trophozoites in the intestinal tract of the host(Chalmers 2014), and 80-90% of asymptomatic infections contribute cysts to the environment(Othman et al. 2020). An infected individual can generate and eliminate into the environment approximately 105 environmentally resistant cysts per gram of fecal material(Haque et al. 2003). Cysts can survive outside the body, including in the sewage system, and they can maintain their viability for up to 2 months(Kott and Kott 1967; Burge and Marsh 1978; Stott 2003; Shirley et al. 2019; Shirley et al. 2020), possibly even after conventional treatment is applied to sewage waters(Razzolini et al. 2020).

In Mexico City and its metropolitan area with a population of approximately 20 million, amoebiasis is considered an important public health concern (Shirley et al. 2019), because of its endemicity (Conde-Bonfil and De La Mora-Zerpa 1992) and circannual behavior(Arroyave et al. 1990). The prevalence of amoebiasis and the factors that modulate its seasonal transmission remain however a matter of debate. Factors affecting the prevalence of *Eh* in Mexico City are considered the result of a combination of individual behavior, the provision of basic infrastructure and sanitation(Sisto et al. 2017), and socio-economic determinants such as poverty and inequality (Cifuentes et al. 2002; Baeza et al. 2018). Less is known about the role of rainfall and the effect on the combined wastewater and stormwater system on driving patterns of disease dynamics. This combined sewage and stormwater system often overflows, exposing residents to street flooding events after heavy rainfall (Romero Lankao 2010; Baeza et al. 2018). Yet, the environmental impact of the urban water system on health outcomes is often overlooked (Garcia-Sanchez and Guereca 2019).

It is possible that rainfall, by overwhelming the capacity of the sewer system during rainy days, can affect the intra- and interannual variability of *Eh* transmission and the observed seasonality of the disease incidence rate. If this is the case, we would expect a significant association and explanatory power of incorporating rainfall in epidemiological models for amoebiasis. If the effect of rainfall is not significant, then the circannual behavior of the disease could be explained by the non-linearity of coupling the epidemiological dynamics of *Eh* with the environmental stages of the pathogen, or by other environmental factors. Understanding the links between disease endemicity and rainfall seasonality could help in prevention, given the relative predictability of the rainy season.

We tested the role of rainfall on the environmental transmission of Eh. in Mexico City with a process-based model of the population dynamics of *Eh* (Fig. 1). The model represents the different stages of *Eh* in the host population, by dividing it into four classes, for susceptible (*S*), exposed (*E*), infectious (*I*), and recovered (*R*) individuals respectively (Hategekimana et al. 2017). A characteristic of the model is the inclusion of an environmental stage of *Eh* outside the host. We specifically included two environmental reservoirs for the pathogen (Fig. 1): The first reservoir (*C*^*R*^) reflected a man-made stormwater system environment in which the concentration of the pathogen was affected by rainfall as a factor influencing overflow See Section 2. The second reservoir represented an environment in which the concentration of the pathogen was not affected by rainfall, *C*^*D*^ (e.g. *Eh* circulating inside houses and/or in the water supply system). The force of infection, *β*_*t*_, is expected to be affected by the exposure of susceptible humans to the pathogen in both reservoirs.

**Figure 1:**
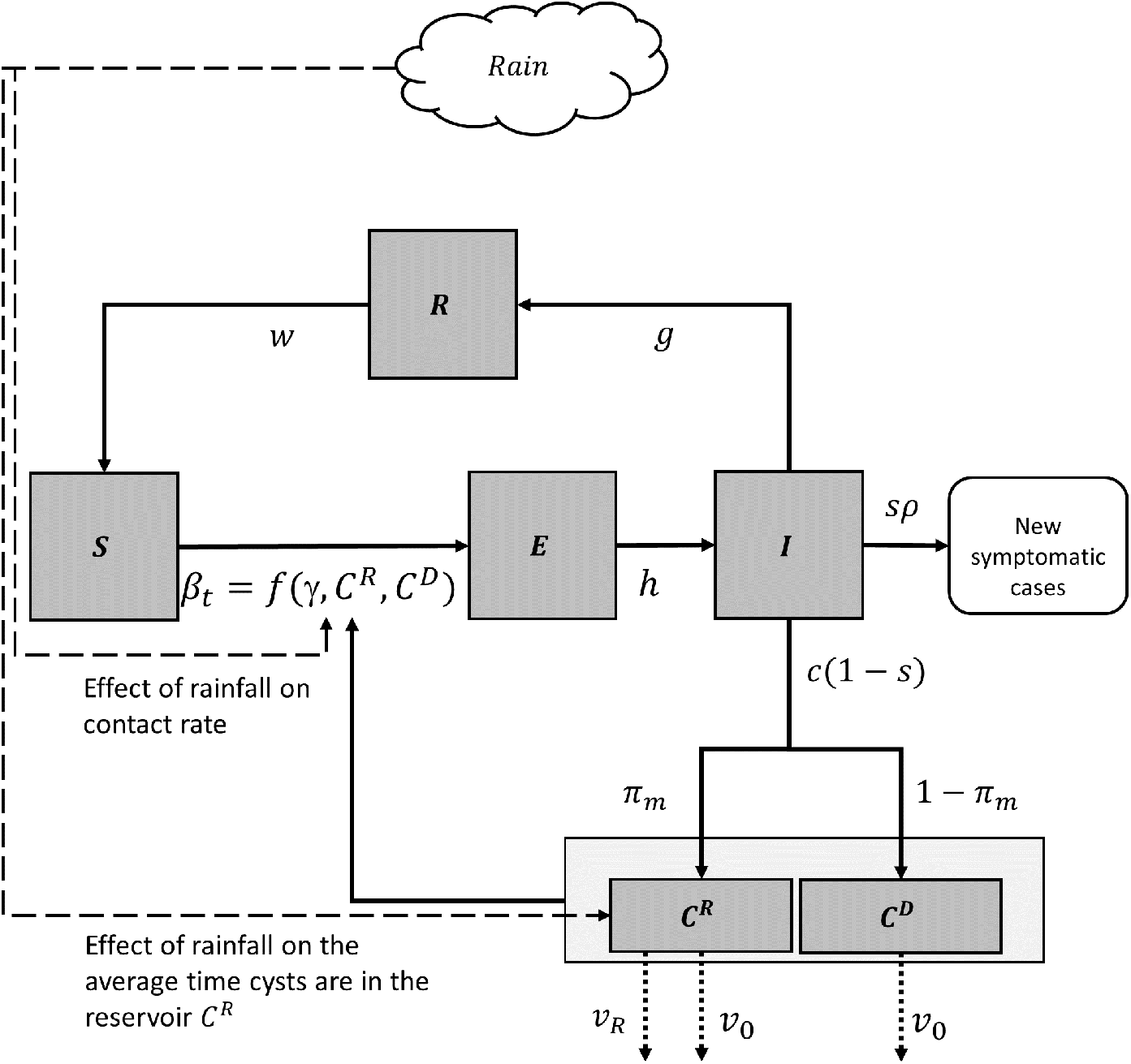
Diagram of the process-based model of amoebiasis.The process-based model is used to simulate the population dynamics of *Entamoeba histolytica* (*Eh*) in Mexico City to test the importance of rainfall on the seasonality of amoebiasis cases. In the model the host population is divided in four classes: susceptible (*S*), exposed (*E*), infected (*I*), and recovered (*R*). The model includes two environmental reservoirs of the cysts of *Eh*. Susceptible individuals become exposed at a rate proportional to the infection rate *β*_*t*_. The infection rate is a function of rainfall. Exposed individuals become infected at rate *h*. New symptomatic infections (*sI*) are reported as cases of amoebiasis, with a reporting rate *ρ*. Infected individual recover at rate *g* and acquire immunity for an average period of 1*/w* years. The concentration of cysts in the rainfall-driven reservoir *C*^*R*^ and in reservoir *C*^*D*^ is affected by the supply of new cysts from asymptomatic individuals *c*(1*− s*). The average time of cysts is the environmental reservoir is 1*/ν*_0_. In the case of the reservoir *C*^*D*^, rainfall also affects the average time cysts are in the reservoir *C*^*R*^ (parameter *ν*_*R*_).

## 2 Methods

### 2.1 Data

#### 2.1.1 Study area and population data

We estimated the model using data from six municipalities in Mexico City that encompassed more than 4.2 million residents. The municipalities are Alvaro Obregon, Benito Juarez, Iztapalapa, Milpa Alta, Tlalpan, and Venustiano Carranza (Fig. 2). These municipalities are located in two zones of the city that differ in biophysical and socio-economic conditions: Tlalpan, Milpa Alta, and the south side of Alvaro Obregon are municipalities located in the “highlands” of Mexico City and that showed the highest proportion of residents without access to infrastructure for drainage and sewer services and contributed the most to the expansion of the urban area. The lack of water-related infrastructure is considered the main source of hydrological vulnerability (Eakin et al. 2016). Previous studies have shown a higher rate of gastrointestinal disease incidence is found in neighborhoods with less access to water-related infrastructure(Baeza et al. 2018).

**Figure 2:**
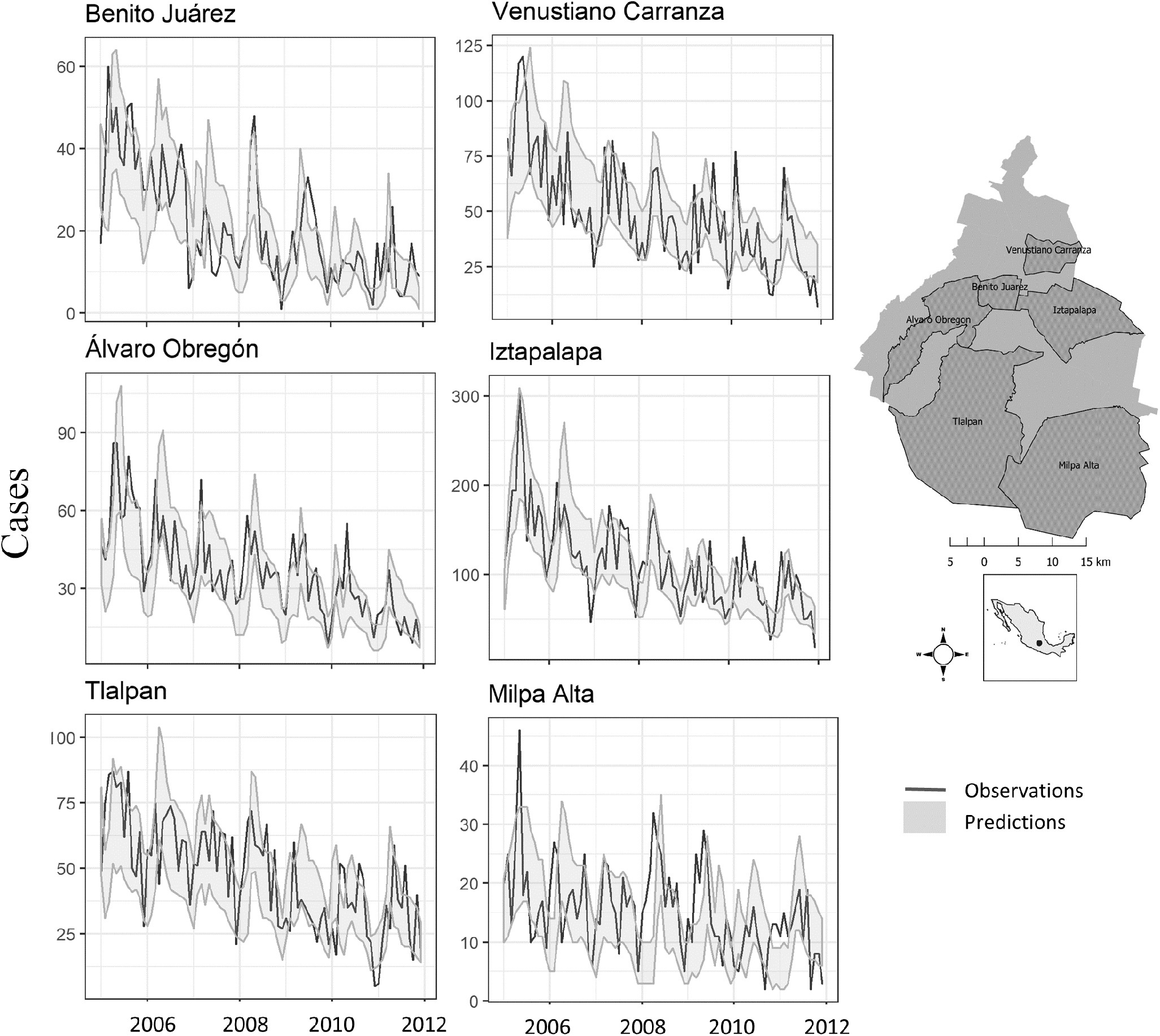
Reported and simulated cases of amebiasis in Mexico City.Using the process-based model, we estimated the effect of rainfall on the population dynamics of *Entamoeba histolytica* in six municipalities of Mexico City, highlighted in dark grey in the Mexico City map to the right of the figure. The six plots to the left of the map show the observations of cases of amoebiasis (black lines) recorded in the public clinics and health centers within of the municipalities between 2005 and 2012. The light grey area is the envelop representing 90

Iztapalapa, Benito Juarez, and Venustiano Carranza and the north side of Alvaro Obregon are municipalities located in the lowlands of Mexico City. These municipalities are located near the ancient Texcoco lake and have been historically vulnerable to flooding. The majority of the houses in these municipalities, unlike those of the highlands, are connected to the sewer system and potable water supply. These municipalities reported the greatest number of street ponding events between 2000 and 2010 (Baeza et al. 2018). Population records from 2010 (*P*_2010_), birth (*b*), and death rates (*d*) were obtained for each municipality from the National Census of 2010 (INEGI 2011). To obtain the population of 2005 (*P*_2005_), we extrapolated backward the population using *P*_2010_ and assuming an exponential growth. Formally,

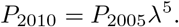

Using the census data, we calculated the growth rate *λ* as:

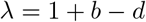

By rearranging it, we obtained

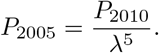

#### 2.1.2 Public records of amoebiasis

We compiled records of confirmed cases of amoebiasis from all clinics in the six municipalities from 2005 to 2011 (Fig. S1). The records were obtained from the Secretaria de Salud de la Ciudad de Mexico, and they consist of confirmed cases of amoebiasis recorded during visits of symptomatic patients to a clinic to which the patient was assigned according to the location of residency. We constructed a single monthly time series of cases by adding all cases from all clinics and hospitals within each municipality recorded each month.

#### 2.1.3 Rainfall records

We obtained records of accumulated daily precipitation [mm/day] for the six municipalities for the same period for which we were able to compile the epidemiological time series (2005-2011). The rainfall data were collected by the Sistema de Aguas de la Ciudad de Mexico (SACMEX) from stations located in strategic places of the city to monitor the capacity of the combined sewer-storm system. From daily records, we constructed an accumulated monthly time series to match the temporal resolution of the disease case data (Fig 2a and Fig S3).

### 2.2 The model

#### 2.2.1 Stages of Entamoeba histolytica in the host population

The population dynamics of the disease are given by the following set of differential equations which divide the population of each municipality into four classes, for susceptible *S*, exposed *E*, infected *I*, and recovered *R* individuals:

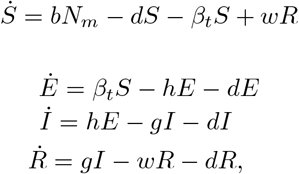

where the total population in municipality *m* is:

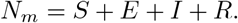

The variable *β*_*t*_ represents the rate of infection, the number of new infections per unit of time. Parameters *b* and *d* are the host’s birth and death rates, and 1*/h* is the mean incubation period from the ingestion of cysts to the development of a trophozoite infection. Thus, *hE*_*t*_ represents the flow of exposed individuals that become infected. Parameter *g* denotes the recovery rate, and parameter *w*, the rate at which recovered individuals lose their immunity (Kantor et al. 2018).

#### 2.2.2 Environmental subcomponent

The model includes two reservoirs to represent the changes over time in the concentration of the parasite in the environment outside the host. The reservoir *C*^*R*^ represents the concentration of pathogens in the environment affected by rainfall, and the reservoir *C*^*D*^, the concentration of the pathogen in an environment not affected by rainfall. Formally, the dynamics of cyst populations are described by the following two equations:

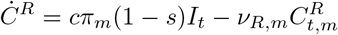

and

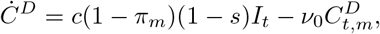

where the state variables 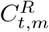 and 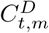 are the respective concentration of cysts in the two reservoirs in municipality *m* at time *t*. Parameter *s* is the proportion of symptomatic infections, and parameter *c* is a conversion parameter linking the number of cysts produced by an infected individual and the concentration of pathogens in the reservoir. Asymptomatic individuals, (1 *− s*)*I*_*t*_, contribute respectively *cπ*_*m*_ and *c*(1 *− π*_*m*_) cysts to reservoirs 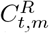 and 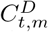 Parameter 1*/ν*_0_ is the average time cysts are existing in the environment. We included in the model the possibility of rainfall also influencing cyst average time in the environment. This effect was captured by parameter *ν*_*r*_. Formally, the rate of change in cyst concentration in the rainfall-driven reservoir is given by the following equation:

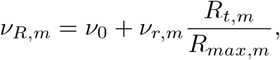

Where the term 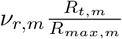 incorporates the hypothesis that rainfall influences the average time cysts are in the reservoir ĈR. *R*_*max,m*_ is the maximum rainfall observed in municipality *m* (Lemaitre et al. 2019).

The force of infection *β*_*t*_ is defined by considering that susceptible individuals are exposed to pathogens in both the reservoir affected by rainfall 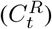 and that not influenced by rainfall 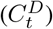. Formally:

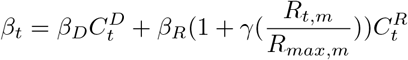

where *β*_*D*_ and *β*_*R*_ are the per-person rate of exposure per dose of pathogen concentration in the reservoirs. Parameter *γ* reflects the effect of rainfall on the contact rate between 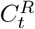 and susceptible individuals, therefore the enhanced exposure to pathogens for every mm of rainfall. Susceptible individuals can also get infected by being in contact with cysts in the reservoir 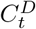, Given the noise in the effects of rainfall on the contact rate, a random effect on the force of infection was included, such that:

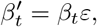

where *ε* represents environmental (dynamical) noise specified with a negative binomial distribution, *ε* = *NegativeBinomial*(0, *σ*).

#### 2.3 Parameter Inference

Since we do not observe the whole system and only monthly incidence is reported by public health centers, we used a partially observed multi-state system. Using the package Partially-observed Monte Carlo Process (POMP), models specify the dynamics of a set state variables that change over time according to some deterministic rules and stochastic uncertainty, with observations for only one or some of these state variables (Ionides et al. 2006; King et al. 2016).

The method consists of maximizing the likelihood estimator of a model to find the best set of estimated parameters that generate the maximum agreement between model simulations and data. The algorithm uses simulation-based methods that treat each simulation (with different combinations of parameters) of the model as a particle. Each particle is evaluated and filtered based on their likelihood estimation, then a new set of particles are created from the ones with a higher likelihood. By an iterated filtering algorithm (MIF), the “best” particles, according to their likelihood value, are selected. These selected particles have a slightly different set of parameter values, to allow exploration of the likelihood surface(King et al. 2016; King et al. 2018).

The model includes a measurement component that links the unobserved state variables to an observed quantity. In our model, the unobserved state is the total number of newly-symptomatic infections, and the observed quantity is the total monthly cases of amoebiasis reported to the public clinics and hospitals in Mexico City, *a*_*t*_ (Fig. 1). New symptomatic infections are reported to clinics in municipality m at a rate *shE*_*t,m*_, where s is the proportion of new symptomatic infections. To link these two states, we defined *E*[*a*_*t,m*_ |*shE*_*t,m*_] as the expected number of amoebiasis cases at time *t*, given the model’s new incidence levels, such that

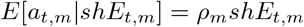

Where parameter *ρ*_*m*_ is the reporting rate. The expected number of amoebiasis cases is simulated as a random process governed by a negative binomial distribution.

### 2.4 Parameter Estimation and likelihood profiles

We first estimated the model parameters for the municipality with the highest population and the highest number of reported cases (Iztapalapa, 1.8 million persons; Table S1). We then used the values of the estimated epidemiological parameters (Table S2) as fixed parameters and the rest of the parameters (those related to the effect of rainfall and the rate of change in the environmental reservoirs) as initial conditions for the other 5 municipalities (Table S2). Using the best-estimated parameters, we simulated the models 10,000 times in each municipality to obtain confidence intervals around the trajectory of the simulated reported cases.

We constructed likelihood profiles for the effect of rainfall on the contact rate ?? and the effect of rainfall on cysts viability *ν*_*R*_. These profiles were calculated by searching the maximum likelihood estimator one at a time for a fixed value of *γ* and *ν*_*R*_, then varied the parameter values systematically near the global equilibrium. If a model with positive values of *γ* and *ν*_*R*_ is significantly “better” than a model estimated for where *γ* = 0 or *ν*_*r*_ = 0, in explaining the data, we can infer using the likelihood estimation and likelihood profiles that rainfall is an important factor in the population dynamics of *Eh*.

#### 2.4.1 Calculating the contribution of rainfall to *Eh* transmission

New infections are all the susceptible individuals that came in contact with cysts of *Eh*. This phenomenon was formalized in the model by the rate of infection, *βS*_*t*_. By integrating the rate of infection over the observational period of our data, we obtained the total number of new infections, *B*. This quantity can be calculated using best-estimated parameters as:

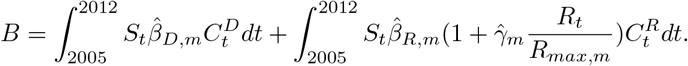

Because the new infections from exposure to the pathogen in the reservoir associated with rainfall can also be calculated, using

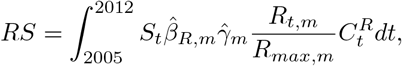

the contribution of rainfall can be estimated as the ratio

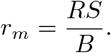

Results of the calculation of *r*_*m*_ are presented in Table S1.

## 3 Results

### 3.1 Seasonality of amoebiasis and rainfall

The cases of Amoebiasis in the six municipalities have a recurrent annual pattern, with the peak of the season occurring between March and May. Between January and the peak of the season, cases steadily increased. A second, but smaller peak occurred between October and November. The most notable feature of the seasonality was the sharp decay in cases observed during December (Fig. 2 and Fig. S1). This decay in cases was observed in all six municipalities. Rainfall data also showed a strong seasonal pattern, where the peak of the rainy season occurred between July, August, and September. In the southmost municipalities (Benito Juarez, Iztapalapa, and Venustiano Carranza), rainfall events exhibited a peak early in the season, whereas those in the north typically peaked later in the year (Milpa Alta, Tlalpan, and Alvaro Obregon). Rainfall was almost completely absent in January, February, March, and December in all municipalities.

### 3.2 Amoebiasis incidence in Mexico City

Iztapalapa was the municipality with the largest number of cases of amoebiasis recorded during this period, accounting for 40% of cases within the 6 municipalities (9433, Table S1 and Fig. S3). The municipality with the least number of cases was Milpa Alta with 1290, corresponding to 5% of the total (Fig. S5). However, Milpa Alta, along with Venustiano Carranza, had the highest incidence risk, with 9.4 and 9.6 cases per 1000 persons, respectively. The number of amoebiasis cases between 2005 and 2011 decreased in all six municipalities (Fig. 1 and Fig. S4, Table S1, and Fig. S5). In Iztapalapa, the number of reported cases dropped from 2112 in 2005 to 881 in 2011 (Fig. 2 and Fig. S3), and in Milpa Alta cases decayed from 233 in 2005 to 135 in 2011.

### 3.3 The contribution of rainfall to amoebiasis transmission

The likelihood profile of the parameter controlling the effect of rainfall intensity on the contact rate (Fig. 3a) showed that the transmission associated with exposure after a rainfall event was significant. Similarly, the profile for the parameter controlling the effect of rainfall on the viability of cysts (Fig. 3b) was significantly different from zero, indicating that rainfall affected the time cysts were present in the man-made reservoir. This likelihood profile analysis indicates that the effect of rainfall was significant in explaining the temporal dynamics of the disease in the city.

**Figure 3:**
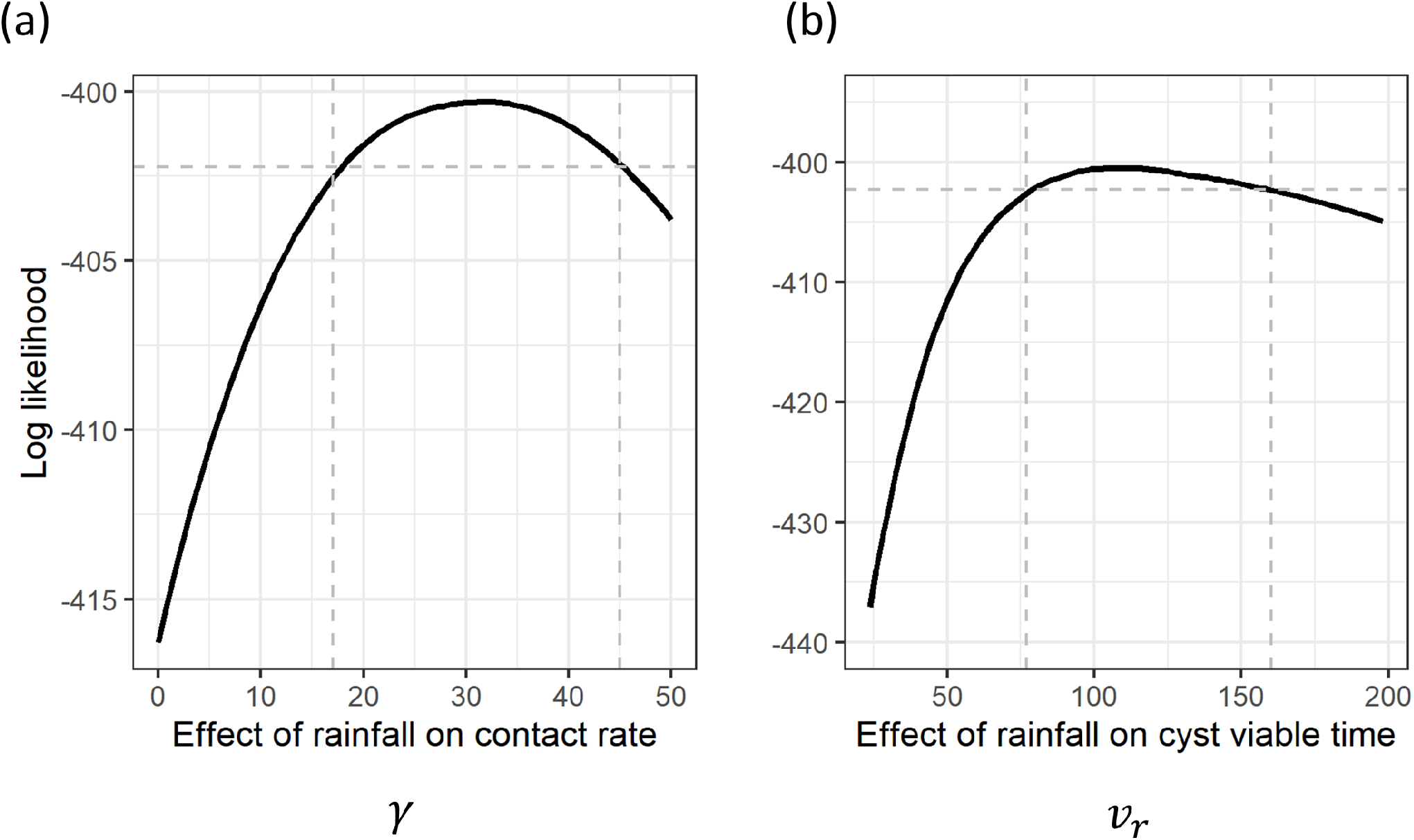
Likelihood profiles of parameters related to the effect of rainfall on the population dynamics of Entamoeba histolytica. Panel (a) shows the maximum likelihood values in a range of values of the parameter controlling the effect of rainfall on the contact rate between susceptible hosts and cysts of Entamoeba histolytica from the reservoir affected by rainfall. Panel (b) shows the maximum likelihood for range of values of the parameter connecting the rainfall to the time cysts are in the reservoir affected by rainfall. The vertical dashed lines represent the 95

Results from 10,000 simulations of the model using the MLE parameters showed that the proportion of the population infected with *Eh* was around 10% in all municipalities in 2005 and decreased to 2 % by 2011 (Fig. S4). The estimation of the model in the six municipalities indicated that the reporting rate of symptomatic infections was around 4% (Table S3). The simulations with the best-estimated parameters showed that the decrease in cases reported between 2005 and 2011 was associated with a decline in the concentration of pathogens in the reservoir not affected by rainfall (Fig. S4).

Using the MLE parameters, we inferred the contribution of rainfall on the transmission by calculating the proportion of the total new infections associated with the rainfall-driven reservoir (*r*_*m*_; See Section 2). The calculations of r_m for the six municipalities indicated that on average more than 90% of new infections resulted from the exposure of susceptible individuals to the pathogens in the reservoir affected by rainfall (Table S1). The municipality with the highest percentage of cases associated with rainfall was Benito Juarez, followed by Iztapalapa and Alvaro Obregon. The municipalities with the least number of cases associated with rainfall were Milpa Alta and Venustiano Carranza.

## 4 Discussion

Our model mechanistically isolates the effects of climate forcing to explain the seasonal dynamics of amoebiasis in Mexico City. Combining epidemiological and rainfall time series data with mathematical modeling and statistical inference, we demonstrated that rainfall had a significant effect on the inter- and intra-annual variability of amoebiasis. These results imply that the chronic hydrological vulnerability of Mexico City (Ezcurra et al. 1999; Romero Lankao 2010; Sosa-Rodriguez 2010) is an important determinant of health at the regional scale. New infections related to increasing the contact rate during the rainfall season accounted for up to 95% of all infections. Our estimations indicate that 10% of the population could have been infected in 2005. The likelihood profiles also indicated that a model that included the effect of rainfall on the average time *Eh* spent in the environment was significantly better than one which did not incorporate this effect.

By including the effect of environmental variability, our model explained why the peak of transmission occurred early in the rainy season. In the model, these factors were considered in the average time the environmental stage of *Eh* remained viable(Kott and Kott 1967). The average timing of *Eh* is often considered to arise from the intrinsic viability of cysts in the environment and the wastewater management and treatments (Kott and Kott 1967; Speich et al. 2016). By including the effect of rainfall with v_r, the concentration of pathogens in the environment was reduced after recurrent days with precipitation. The result was the reduction in the risk transfer to the population at the peak of the rainfall season. In contrast, during the dry season, the sewer system carried water with a higher concentration of pathogens than during the wet season. Thus, rainfall events early in the season could be most important in driving transmission due to a combination of high pathogen concentration and high contact rates.

Results from the parameter estimation and the simulations of the unobserved states showed that the risk to *Eh* exposure in the reservoir not affected by rainfall decreased during the observational period. This decrease in risk helps to explain the downtrend in amoebiasis cases observed in the hospital records. Presumably, this downtrend in the data was the result of improvements in sanitation, wastewater treatments, and behavioral interventions. Our estimations also indicated that the risk associated with the reservoir affected by rainfall has not declined. Because of the conceptualization of a reservoir affected by rainfall reflected man-made structures, these results suggest that the risk associated with the infrastructure used for climate and water management is an important determinant of health. Moreover, the dominance of the seasonal climate-driven reservoir on the dynamics of *Eh* suggests that Mexico City residents could be exposed to chemical- or biohazards whose frequency of exposure is associated with the city’s rainfall season.

The impact of rainfall as an external forcing of health risk in Mexico City should supports calls for more integrated health and water management planning as part of climate adaptation policy. This is particularly the case when climatic patterns are highly seasonal, and infrastructure is inadequate to manage existing water and sanitation demands and to confront climate change. The results presented underscore the importance of complementing behavioral or engineering interventions with environmental ones designed to halt transmission. For instance, such efforts could incentivize awareness of the risk posed by rainfall in educational campaigns early in the rainy season. This study contributes to a body of knowledge on how climate modulates the population dynamics of pathogens in megacities (Reiner et al. 2012; Martinez et al. 2016; Santos-Vega et al. 2016a). The approach used herewith should be extended to other protozoan parasites with environmental resistance stages such as Giardia. It also should continue for other cities affected heavily by climate forcing. A global picture of the future environmental health risk posed by climate change is important to elucidate and differentiate individual vs. public responsibilities and would help to define more accurate environmental health interventions.

## 5. Acknowledgments

We would like to thank the National Institute of Statistics and Geography (INEGI), the Secretary of Health of Mexico City, and the Mexico City water operator (SACMEX) for the data provided. We thank Dr. Mercedes Pascual and Brieta Fass for their valuable comments and editions of early versions of this manuscript. This research was funded by the National Science Foundation (Grant No. 1414052) and the Institute for Social Science Research (ISSR) at ASU. Any results, errors, or interpretations presented in the manuscript are the responsibilities of the authors and not of the funding agencies.

