## Supplementary information for "The unapparent effect of climate forcing on urban amoebiasis in Mexico City"

### **Supplemental files**

Table S1: Contribution of rainfall to new infections. The table shows the population in each municipality, the number of amoebiasis cases and the incidence rate between 2005 and 2011. The table also shows the mean value of the ratio $r_m$, which is the calculation of the proportion of new infections attributed to the exposure of susceptible individuals to *Eh* from the rainfall-driven reservoir. The mean value was obtained after 10000 simulations of the mechanistic model using best-estimated parameters.

| **Municipality** | **Population 2010** | **Total cases** | **Incidence rate** | $r_m$ |
|---|---|---|---|---|
| Benito Juarez | 417,416 | 1,747 | 4.2 | 97% |
| Iztapalapa | 1,827,868 | 9,433 | 5.2 | 96% |
| Alvaro Obregon | 749,982 | 3,090 | 4.1 | 95% |
| Tlalplan | 677,104 | 3,951 | 5.8 | 95% |
| Milpa Alta | 137,927 | 1,290 | 9.4 | 93% |
| Venustiano Carranza | 427,263 | 4,088 | 9.6 | 84% |

Table S2: Regional parameters. The table described the parameters of the *Eh* model estimated in Iztapalapa and maintained fix for the full city. Parameters with reference were maintained constant and not subjected to estimation.

| **Name** | **Symbol** | **Value** | **Reference** |
|---|---|---|---|
| Incubation period | $h$ | 1/14 days | (Gerba 2015, Hategekimana et al 2017) |
| Birth rate | $b$ | 0.0183 | INEGI 2011 |
| Death rate | $d$ | 0.0058 | INEGI 2011 |
| Average length of an infection | $g$ | 5 months | Estimated |
| Immunity | $w$ | 1.08 years | Estimated |
| The proportion of symptomatic infections | $s$ | 0.096 | Estimated |
| Cyst production per infection | $c$ | 0.94 | Estimated |

Table S3: Parameters estimated by municipality. The table shows the description and symbol of the parameters of the amoebiasis model and the best estimated parameter values for each of the six municipalities in Mexico City.

| Description | Symbol | Best estimated parameters by Municipality | | | | | |
|---|---|---|---|---|---|---|---|
| | | Iztapalapa | Milpa Alta | Alvaro Obregon | Benito Juarez | Tlalplan | Venustiano Carranza |
| Proportion of cysts released into $C_m^R$ | $\pi_m$ | 0.227 | 0.24 | 0.23 | 0.22 | 0.2431 | 0.2271 |
| Cyst viability | $\nu_{0,m}$ | 0.71 | 1.5 | 0.73 | 0.74 | 0.718 | 0.67 |
| Rainfall effect on cyst viability | $\nu_{R,m}$ | 260 | 130 | 140 | 170 | 144 | 175 |
| Per-person rate of infection associated with reservoir $C_{D_0,m}$ | $\beta_{\mathrm{D},m}$ | 6.87 | 16.5 | 5 | 2.6 | 6.87 | 8 |
| Effect of rainfall on the contact rate between pathogens in $C_m^R$ and susceptible individuals | $\gamma_m$ | 200 | 140 | 143 | 320 | 120 | 55 |
| Per-person rate of infection associated with $C_m^R$ | $\beta_{R,m}$ | 4.8 | 6 | 4.4 | 2.7 | 5 | 11.2 |
| Initial proportion of pop. susceptible | $S_0$ | 0.62 | 0.55 | 0.5 | 0.5 | 0.3 | 0.45 |
| Initial proportion of pop. exposed | $E_0$ | 0.00492 | 0.00425 | 0.0048 | 0.0049 | 0.004 | 0.01 |
| Initial proportion of pop. Infectious | $I_0$ | 0.101 | 0.103 | 0.108 | 0.105 | 0.101 | 0.103 |
| Initial proportion of pop. immune or recovered | $R_0$ | 0.27408 | 0.34275 | 0.3872 | 0.3901 | 0.595 | 0.437 |
| Initial concentration of cysts in $C_m^R$ | $C_{0,m}^R$ | 0.0005 | 0.0002 | 0.0002 | 0.0003 | 0.0004 | 0.0003 |
| Initial concentration of cysts in $C_m^D$ | $C_{0,m}^D$ | 0.027 | 0.025 | 0.025 | 0.07 | 0.04 | 0.09 |
| Reporting rate | $\rho_m$ | 0.046 | 0.039 | 0.037 | 0.039 | 0.0398 | 0.038 |

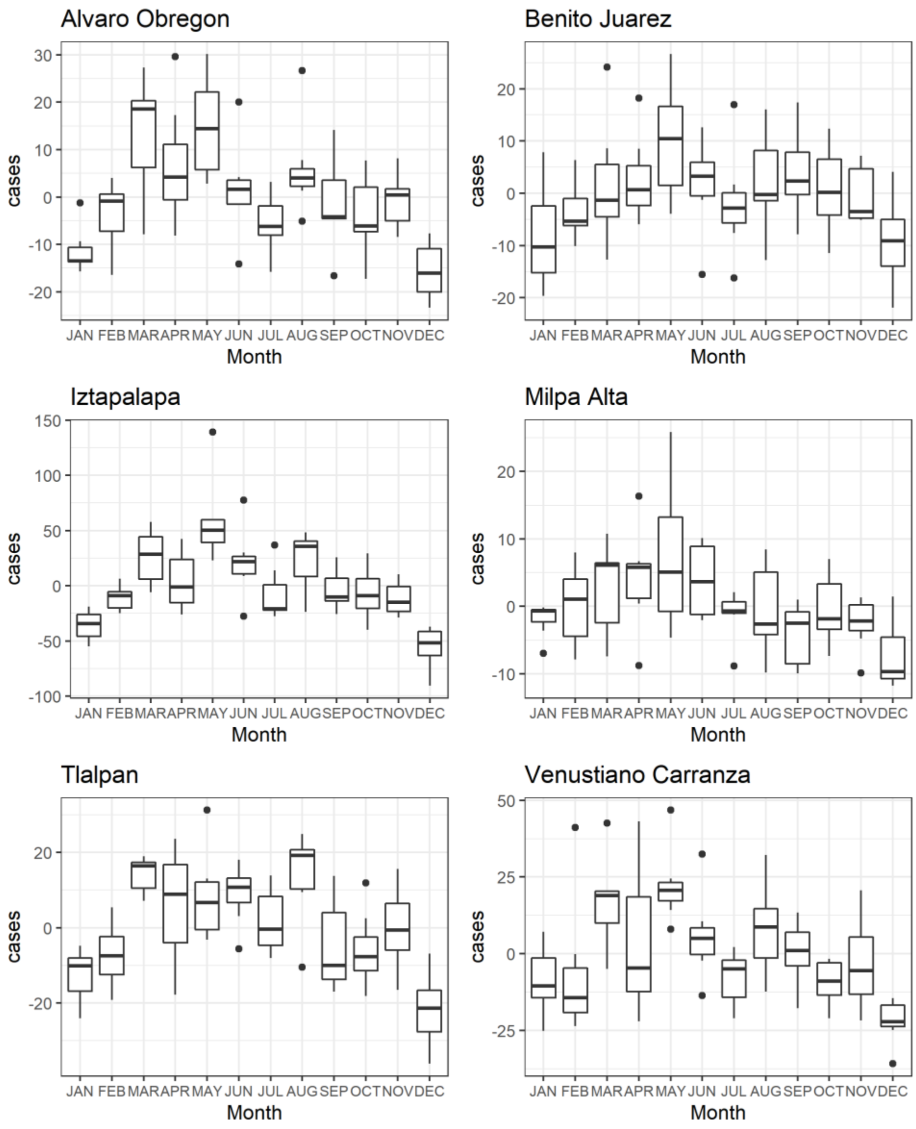


Figure S1: The seasonality of amoebiasis in six municipalities of Mexico City from 2005 to 2011. Each boxplot shows the monthly cases as deviation from the mean (horizonal dark lines), standard deviation (box), standard error (vertical lines), and outliers (dots).

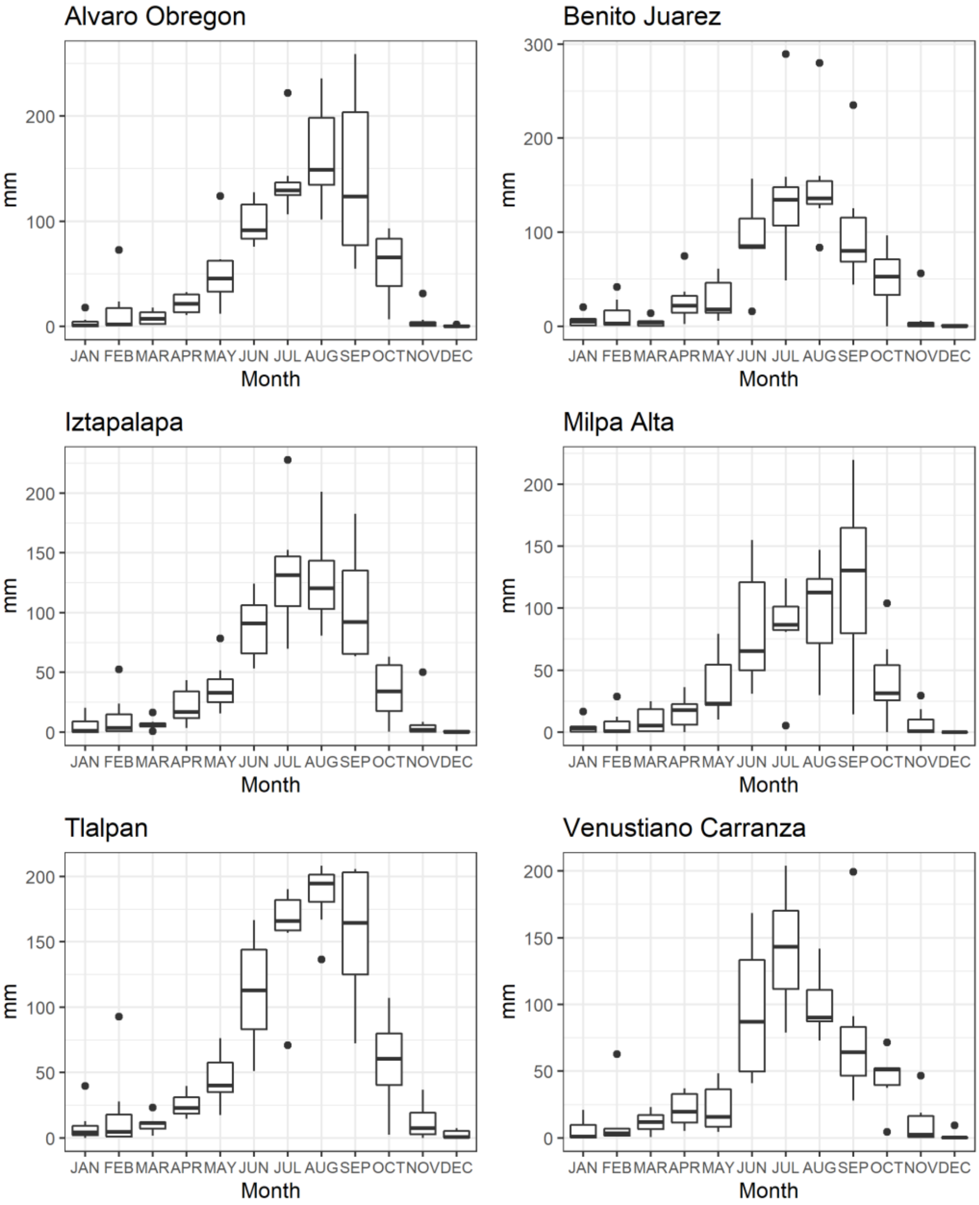


Figure S2: Monthly rainfall seasonality in six municipalities of Mexico City. The boxplots show the average rainfall in a month (mm; horizonal dark lines), standard deviation (dashed bars), standard error (horizonal box edges), and outliers (shallow dots) from 2005 to 2012.

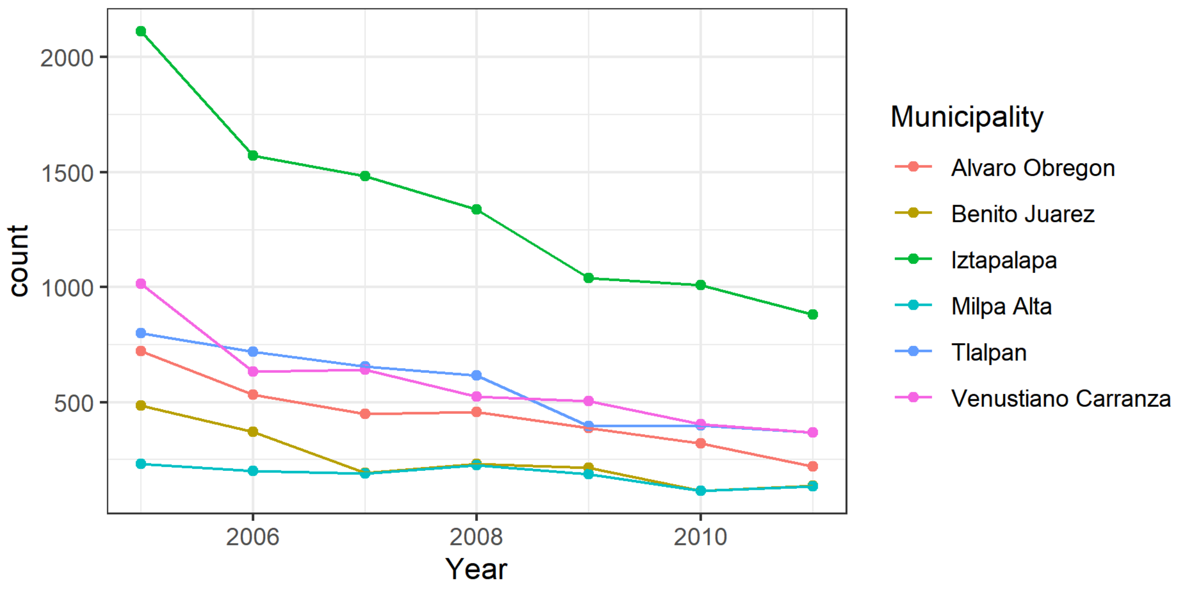


Figure S3: Annual amoebiasis cases. The figure shows the decreasing trend in annual cases of amebiasis in 6 municipalities of Mexico City between 2005 and 2011.

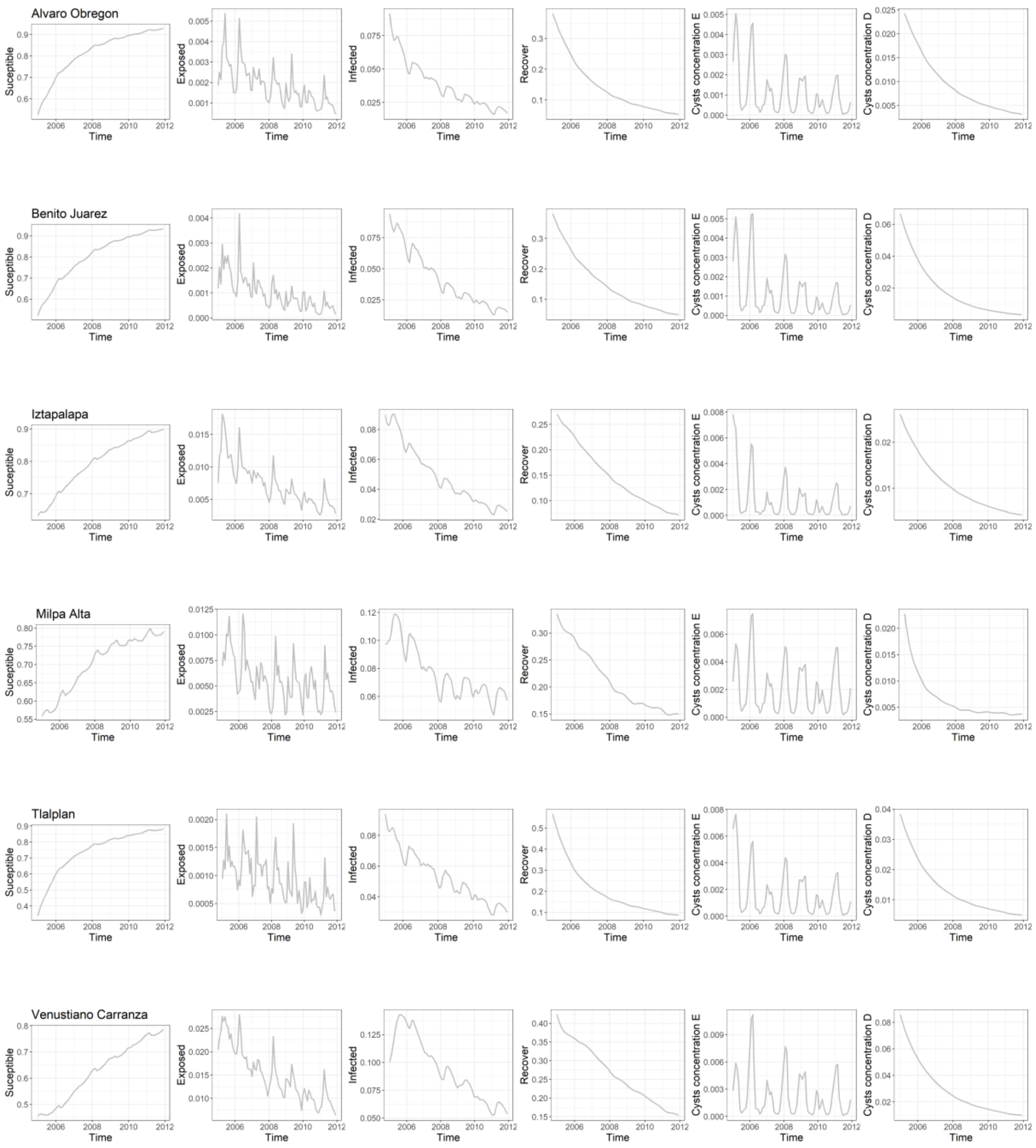


Figure S4: The six unobserved states of the population dynamics of *Entamoeba histolytica* in six municipalities in Mexico City. Each plot shows the dynamics of the state of a single simulation using best-estimated parameters.
